# Rab5-mediated Yolk Cell Endocytosis modulates Zebrafish Epiboly Biomechanics and Tissue Movements

**DOI:** 10.1101/097212

**Authors:** Maria Marsal, Amayra Hernández-Vega, Philippe-Alexandre Pouille, Enrique Martin-Blanco

**Affiliations:** Instituto de Biología Molecular de Barcelona, Consejo Superior de Investigaciones Científicas Parc Cientific de Barcelona, Baldiri Reixac 10, 08028 Barcelona, Spain

## Abstract

Morphogenetic processes demand the coordinated allocation of cells and tissues to their final destination in a spatio-temporal controlled way. Identifying how these morphogenetic movements are directed and implemented is essential for understanding morphogenesis. Topographical and scalar differences in adhesion and contractility within and between cells are essential, yet, the role that membrane remodeling may play remains less clear. To clarify how surface turnover and dynamics may modulate tissue arrangements we studied epiboly in the zebrafish. During epiboly the blastoderm expands as a result of an asymmetry of mechanical tension along the embryo surface. In this scenario, we found that the membrane removal by macropinocytosis of the external yolk cell syncytial layer (E-YSL) ahead of the blastoderm is key for epiboly progression In early zebrafish embryos, the activity of the GTPase Rab5ab was essential for endocytosis, and interference in its expression exclusively in the yolk cell resulted in the reduction of yolk cell actomyosin contractility, the disruption of cortical and internal yolk flows, a disequilibrium in force balance and as a result epiboly impairment. We conclude that regulated membrane remodeling is crucial for directing cell and tissue mechanics and coordinating morphogenetic movements during epiboly.

## INTRODUCTION

Membrane turnover can be clathrin-dependent, caveolae-mediated or via macropinocytosis. Clathrin-mediated endocytosis (CME), a mechanism for controlled cargo uptake, is the most general membrane internalization route. CME is characterized by the formation of clathrin-coated vesicles (CCV) and the selective internalization of cell-surface components and extracellular macromolecules (for review see [1]). CME depends on Dynamin activity for vesicle budding and scission, which leads to trafficking through the endocytic pathway upon clathrin coat disassembling. Alternatively, caveolae (which incorporate caveolin) are also widely employed by cells for membrane removal. They originate in the Golgi apparatus, associate to cholesterol pools and are involved in the internalization of specific markers and perhaps in transcytosis. Last, large-scale membrane internalization can also take place by actin-dependent macropinocytosis, which usually occurs within highly buckled regions of the plasma membrane [2-4].

Endocytosis and recycling have been shown to affect cell surface areas [5, 6] and tension and contraction [7, 8] at different levels, from cellular maturation (e.g. dendritic arborization in the *Drosophila* larvae [9]) to morphogenesis (reviewed in [10]). Cellularization and dorsal closure in *Drosophila* and neurulation and apical constriction of bottle cells during gastrulation in *Xenopus laevis* are just some examples of processes demanding a tight control of membrane dynamics [6, 11, 12].

Rab5, a member of the Rab guanosine triphosphatases (small GTPases) family is implicated in CME [13] and the internalization and merging of vesicles into early endosomes (reviewed in [14]). Rab 5 and its effector rabankyrin 5 are also involved in macropinocytosis [15-17]. Both, Dynamin and Rab5 cooperate downstream of actomyosin contractility to remove excess membrane; e.g. disrupting endocytosis either with dominant-negative Dynamin or Rab5 perturbs neurulation in *Xenopus*, inhibiting the apical constriction of hingepoint cells and triggering neural tube closure defects [6]. In the zebrafish, endocytosis affects Silberblick (slb)/Wnt11 activity and E-cadherin trafficking, both necessary for epiboly [18]. It also affects the ability of cells to aggregate into clusters via E-cadherin, which affects the coordinated movement of the prechordal plate [19]. In the fish embryo, Rab5 also mediates the cohesion of mesodermal and endodermal (mesendodermal) progenitor cells during gastrulation. In summary, the modulation of the balance between endocytosis and recycling has a clear-cut role on the regulation of cellular morphology and tissue deformations.

We aimed to understand the role that membrane remodeling might have during the conserved early morphogenetic movements leading to epiboly in the zebrafish. At the onset of zebrafish gastrulation (sphere stage), a superficial layer of cells, the enveloping layer (EVL) covers a semi-spherical cap of blastomeres sitting on a massive yolk syncytial cell and centered on the animal pole of the embryo. Epiboly consists on the cortical vegetal ward expansion of the EVL, the deep cells (DCs) of the blastoderm and the external layer of the syncytial yolk cell (E-YSL) around the yolk. It ends with the closure of the EVL and the DCs at the vegetal pole [20-22] (Figure 1A).

**Figure 1.**
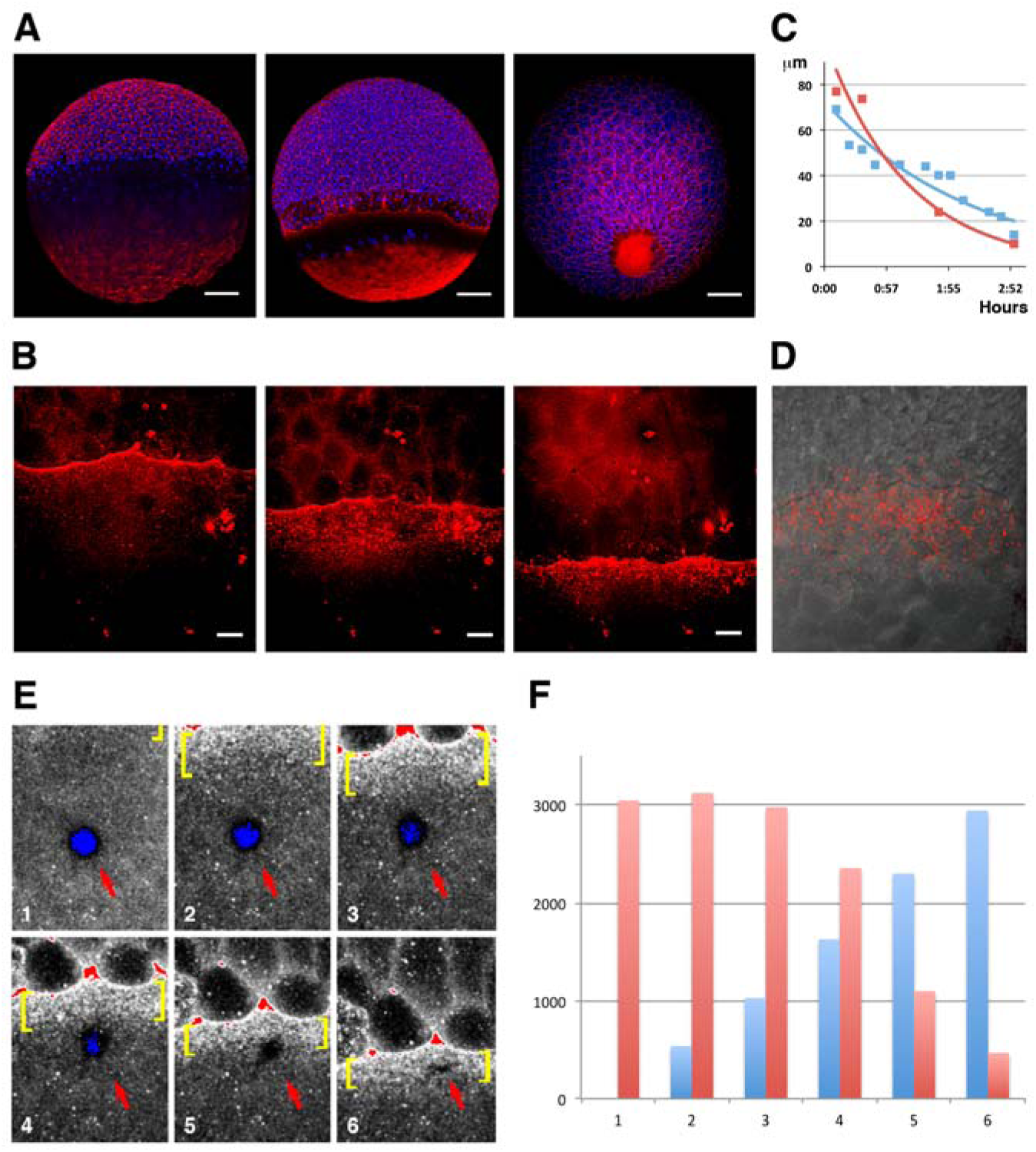
Yolk cell membrane endocytosis at the E-YSL. **A)** Blastoderm expansion during epiboly. At sphere stage epiboly has not yet begun (left). F-actin accumulates at the periphery of all cells as well as in the yolk cell, mainly at the vegetal cap. At 70 % epiboly (middle), the blastoderm has crossed the equator and will decrease its margin until closure. A belt of actin develops at the E-YSL ahead of the EVL and an actin-free zone separates this belt from a vegetal actin-rich patch. At 90 % epiboly (right), the E-YSL and the vegetal actin-rich patch merge. Embryos were stained with phalloidin-TRITC (red) and DAPI (blue). Scale bars 100 μm. **B**) Sequential images of a confocal time-lapse video of a wild type embryo soaked in lectin–TRITC for 5 minutes at sphere stage. The lectin binds to the membrane of both the yolk cell and the EVL cells and gets internalized accumulating in vesicles in the E-YSL just ahead of the EVL margin (from **Movie S1**). Scale bar 25 μm. All confocal images are maximum projections. **C)** Parallel reduction during epiboly progression of the width of the convoluted domain (blue) and of the area undergoing membrane removal (red) of the EYSL. X and Y axes represent hours after 50 % epiboly and width in μm, respectively. **D)** Uptake of fluorescent dextran (red) at the E-YSL just ahead of the EVL margin at 65 % epiboly. **E**) Snapshots of time-lapse images (from **Movie S2**) of a lectin-TRITC soaked embryo showing a circular photobleached area (red arrow) in the yolk cell away from the EVL leading edge. The photobleached membrane is removed and endocytosed only upon its enclosure within the advancing E-YSL (yellow brackets). All confocal images are maximum projections. Scale bar 25 μm. **F**) Membrane internalization dynamics. Histograms depicting the normalized relationship between the photobleached areas (red) and the progression of the leading EVL front (blue) at regular sequential time points (from 1 in **E**). The removal of the photobleached membrane shows a precise linear relationship with the speed of advancement of the leading front.

Epiboly progression entails a coordinated series of cellular events. EVL cells and DCs proliferate and exchange neighbors resulting in EVL expansion and DCs intercalation [7, 23]. At the yolk cell, the membrane of the proximal E-YSL membrane becomes highly buckled [24]. This domain gradually narrows by localized contraction [25]. As the EVL margin advances, myosin and polymerized actin get progressively recruited to the convoluted belt at the animal edge of the E-YSL and to the vegetal cap [23, 26]. At the proximal E-YSL, actin is conscripted within and beneath the highly dynamic buckles [25]. Actin accumulation is accompanied by myosin phosphorylation [26] and, remarkably, the narrowing of the E-YSL occurs in synchrony with cortical retrograde flows of actin and myosin originating at the vegetal pole [27]. Internally and coupled to epiboly progression, the yolk granules sustain stereotyped coordinated movements [25].

It has been suggested that longitudinal and latitudinal tensional forces originating at the E-YSL constitute the major force-generating elements driving epiboly [23, 26, 28, 29] and it It has also been proposed that for the vegetal ward movement of the blastoderm some source of tension must be coupled to the contractile E-YSL. Flow-friction motors could generate a pulling longitudinal force through resistance against the retrograde actomyosin cortical flows in the yolk [27]. Alternatively, a positive vegetal ward oriented latitudinal tension gradient at the yolk cell membrane from the EVL margin could convey the stress originated by the constriction of the actomyosin ring at the EYSL [25].

Yet, whatever the mechanical means, epiboly must overcome the hindrance that the yolk cell membrane poses to its progression. The EVL does not slide over the yolk cell surface and is firmly attached to it [25, 26]. Thus, as it expands, the external yolk cell membrane subsides and it is fully eliminated [7, 23, 28] pointing to membrane removal at the yolk cell as an essential necessary event for epiboly progression [7, 28].

To analyze the role of yolk cell membrane remodeling during epiboly we first spatially and temporally characterized its turnover as epiboly proceeded. We also found that the activity of Rab5ab, a key element of the endosomal route and mediator of macropinocytosis [14], which was previously found to affect gastrulation [30], was essential for the removal of the yolk cell membrane and epiboly movements. Our results differ from previous data gathered by interfering with Dynamin 2 activity (clathrin-mediated endocytosis) [31], which indicated that the removal of the yolk cell membrane was dispensable for epiboly.

We showed that Rab5ab was essential for the recruitment of actin and myosin to the EYSL and for their contractile activities. It is essential to sustain the yolk cell surface tension reinforcing the strength of the E-YSL as a mechanical power source. Altering this biomechanical landscape results in a failure of EVL cells elongation, weakening of internal yolk flows and epiboly’s failure. Altogether, our data show that localized membrane removal in the yolk cell constitutes a necessary step for epiboly progression bridging cellular, geometrical and mechanical constrains.

## RESULTS

### E-YSL membrane dynamics

To study zebrafish embryo membrane turnover we employed fluorophore-conjugated lectins. Lectins bind to glycoproteins and glycolipids and have already been used to follow plasma membrane dynamics in other teleost embryos [23]. Upon soaking the embryos in fluorophore-conjugated lectin-containing media, both, the whole yolk cell and the EVL external membranes were quickly homogeneously decorated. Immediately after, lectin-enriched spots, resembling endocytic vesicles, deposited beneath the yolk cell membrane, accumulating in a circumferential ring ahead of the EVL leading cells (Fig. 1B and **Movie S1**). Membrane removal to the front of the EVL can be observed as early as the sphere stage and co-localized with the convoluted E-YSL, where actin and myosin progressively gather [27]. We observed a tight spatiotemporal correlation between the narrowing of the endocytic belt and the reduction of the width of the EYSL (convoluted yolk cell surface) (Fig. 1C), which suggests an intimate relationship between both events. Remarkably, fluid phase endocytosis, reported by the uptake of fluorescent dextran [30, 32], displayed the same pattern as lectin internalization (Fig. 1D).

To precisely map and characterize yolk cell membrane turnover, we locally marked the membrane by laser photobleaching and followed its dynamics *in vivo*. Membrane photobleached regions away from the EVL edge remained static, indicating the lack of major lateral diffusion within the yolk cell membrane up to the time when the photobleached areas of the yolk cell got embedded in the advancing E-YSL. At this time the tagged membrane subdued and became endocytosed (Fig. 1E and **Movie S2**). The photobleached area linearly reduced its size and was finally eliminated before contacting the EVL margin (Fig. 1F).

The observed dynamics of the yolk cell membrane confirms that the EVL does not slide over the yolk cell [26] and suggests that its progression demands the E-YSL progressive removal in an animal-vegetal direction up to its full disappearance, so that the overall surface of the yolk cell remains constant.

### Rab5ab-mediated endocytosis is required in the yolk cell for epiboly progression

We found that the turnover of the yolk cell membrane is topographically associated with the E-YSL proximal domain. The extensive convolution of this area [25] points to nonclathrin dependent macropinocytosis as the mechanism for its removal. Indeed, abolishing Dynamin-2 expression in the yolk cell just results in a partial inhibition of endocytosis not affecting epiboly progression [31].

To fully block membrane removal we set out to interfere with *rab5* expression, a key element for both vesicle internalization and targeting to early endosomes and for macropinocytosis [14]. To target only the yolk cell without affecting the blastoderm, we performed morpholino (MO) injections into the yolk syncytium (YMOs) after its segregation at the 512-1024 cell-stage. In this way, injected MOs remained confined to the yolk cell and were not mobilized to the rest of the embryo [33]. Yolk injection of mRNAs or fluorescently-tagged MOs at these stages showed restricted expression in the yolk cell (Fig. S1).

In zebrafish, there are five annotated *rab5* genes (*rab5aa*, *rab5ab*, *rab5b*, *rab5c* and *rab5clike*). Of these, *rab5aa*, *rab5ab*, and *rab5c* were known to be ubiquitously expressed, unlike *rab5b*, whose expression was limited to the yolk cell syncytial layer, pronephric duct, and telencephalon [34].

We tested whether inhibition of different *rab5* genes prevented membrane removal by monitoring fluid-phase dextran endocytosis. Three different *rab5ab* morpholinos, Rab5ab YMO1 (ATG), YMO2 (UTR) and YMO3 (UTR-ATG) injected in the yolk caused the same phenotype, while a control mismatched MO or a *rab5c* YMOs [35] did not. Note that MO1 and MO2 had been previously employed to analyze the role of Rab5ab in nodal signaling during gastrulation. In that context, the effect of both MOs on *gsc* expression was fully rescued by co-injection of a *rab5ab* RNA ensuring their specificity [30].

We found that Rab5ab depletion just in the yolk cell led to deficient membrane removal (Fig. 2A and **Movie S3**). Quantitative analysis further showed that the number of internalized dextran-containing vesicles in Rab5ab YMOs was reduced by 84 % (n = 7), while interfering with Dynamin 2 expression in the yolk cell reduced this figure by 48 % [31]. Importantly, yolk cell specific Rab5ab depletion resulted in a strong early epiboly delay and arrest while other gastrulation and morphogenetic movements (invagination, convergence and extension, and somitogenesis) and head and trunk development initiated timely and seemed mostly unaffected (see Fig. 2B and **Movie S4**). Rab5ab YMOs displayed a dose-dependent response. Prior to epiboly no apparent defect was observed at any dosage and phenotypes were manifested from dome stage onwards. At a medium dose (4 ng / embryo), *rab5ab* YMOs domed in a timely manner but immediately slowed down, halting at 70 % epiboly. When control injected embryos reached the shield stage, medium dose Rab5ab YMOs had not progressed beyond 30 % and when the DCs of controls closed the yolk cell plug, Rab5ab YMOs still remained at 60 % epiboly (compare Fig. 2C to 2D; see **Movie S5**). High dose yolk cell-injected embryos (8 ng / embryo) never progressed beyond 50 % epiboly and burst shortly after, (compare Fig. 2C to 2E; see **Movie S6**). Epiboly arrest correlated with a progressive buckling of the DC layer detaching from the YSL, and with a constriction at the yolk surface ahead of the EVL margin. Alongside the epiboly delay, Rab5ab YMOs failed to thin the blastoderm, which retracted animalward.

**Figure 2.**
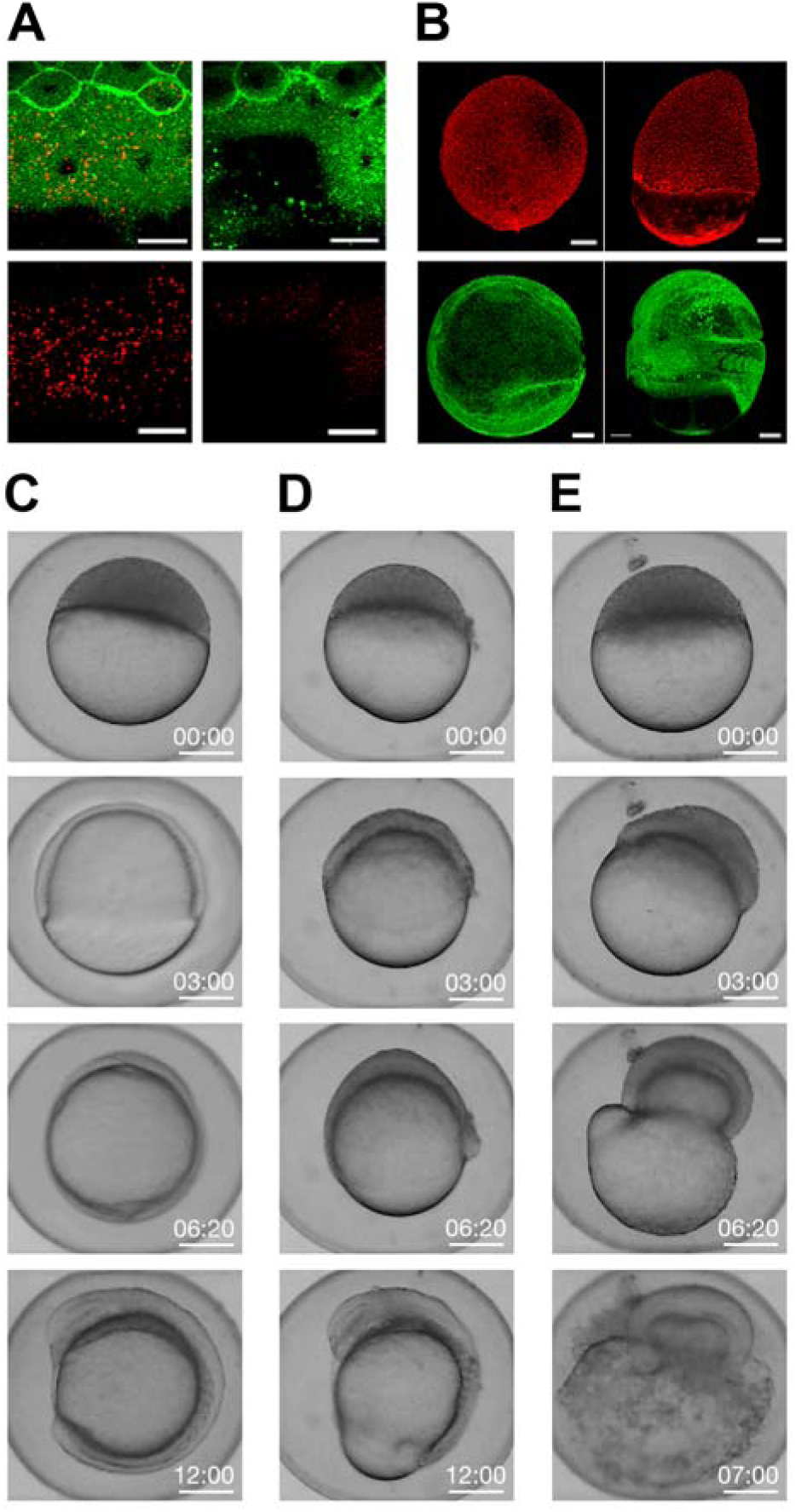
Endocytosis and Epiboly are impaired by Rab5ab depletion. **A**) Rhodamine B-Dextran internalization was reduced at doming stage in membraneGFP (Tg (β-*actin:m-GFP) rab5ab* (right) compared to sibling controls (left) YMOs. Top panels show dextran (red) and membrane staining (green) overlays. Images are confocal maximum projections. Scale bar 25 μm. See also **Movie S3**. **B**) Yolk cell depletion of Rab5ab results in epiboly delay without affecting other gastrulation movements [Control (left) and Rab5ab YMOs (right)]. Top panels show Phalloidin-TRITC (red) staining of a control YMO at the end of epiboly and a sibling medium dose Rab5ab YMOs of the same age. Lateral views. Bottom panels show membrane-GFP (Tg (β-*actin:m-GFP*) YMOs at 1 day post fertilization. Rab5ab YMOs present an open back phenotype but succeed in other gastrulation movements leading to somite formation (see also **Movie S4**). Images are confocal maximum projections. Scale bar 100 μm. **C-E**) YMOs show a dose dependent epiboly delay. Macroscopic bright field images of sibling controls (**C**) and medium (4 ng) (**D**) and high (8 ng) dose (**E**) Rab5ab YMOs (from **Movies S5** and **S6**). Medium and high dose YMOs remained at 70 and 40 % epiboly respectively when control siblings have already closed. Embryos were imaged in their chorion. Scale bar 250 μm.

Altogether, these data indicate that *rab5ab* dependent endocytosis is involved in local yolk cell membrane removal at the E-YSL and that this is necessary to enable epiboly progression.

### Rab5ab activity in the yolk cell affects cortical actomyosin and, non-autonomously, EVL shape and yolk granules dynamics

The epiboly progression failure observed after inhibition of Rab5ab expression in the yolk cell was associated to other cellular and structural phenotypes. While, in principle, these defects could be connected to pleiotropic cell signaling faults, their absence upon interference in the yolk cell of other Rab5 isoforms (e.g. Rab5c, which affects E-Cadherin turnover and epiboly and the movements of the prechordal plate when inhibited in full embryos [18, 35]) supports their potential bonds with Rab5ab-mediated bulk effects on membrane removal.

The local recruitment to the E-YSL of actin and phosphorylated myosin was compromised by middle dose interference in *rab5ab* expression in the yolk cell. The levels of both proteins, detected with phalloidin and anti phospho-myosin antibody respectively, were strongly reduced in Rab5ab YMOs (Fig. 3A and 3B). These reductions correlate with defects in the retrograde cortical myosin flow observed in wild-type animals [27]. Live time-lapse imaging of transgenic Tg (β-*actin:MYL9LGFP*) embryos revealed that the magnitude of the yolk cell cortical myosin flows was decreased and their directionality altered in Rab5ab YMOs (Fig. 3C and **Movie S7**). Moreover, the narrowing of the actin-rich convoluted E-YSL, the major source of force generation during epiboly [25], was also affected. The E-YSL became wider overtime (as quantified from surface projections of membrane-GFP tagged embryos - see Experimental Procedures) (Fig. 3D).

**Figure 3.**
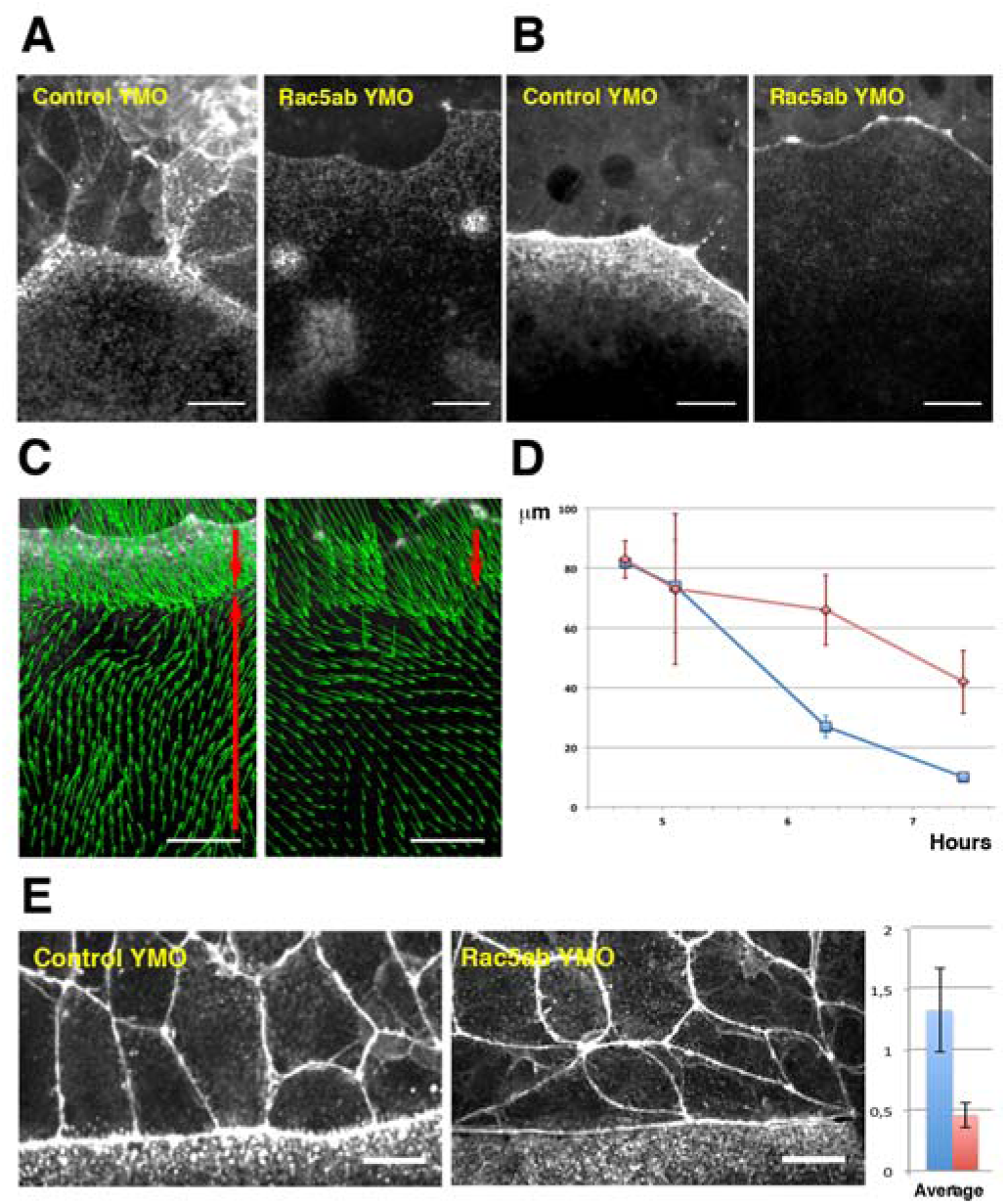
Cytoskeleton dynamics and EVL leading cells shapes are affected by Rab5ab depletion. **A)** Actin fails to accumulate at the E-YSL in Rab5ab YMOs versus controls (C YMO). Time-lapse snapshots of two LifeAct GFP injected sibling embryos. **B)** Myosin fails to accumulate at the E-YSL of Rab5ab YMOs versus controls (C YMO). Time-lapse snapshots of two Myosin-GFP transgenic (Tg (β-*actin:MYL9L-GFP*)) sibling embryos. Notice the delay in the progression of the EVL and the weaker accumulation of actin and myosin in the Rab5ab YMOs. **C**) Myosin cortical retrograde flows. PIV of time-lapse snapshots of Tg (β-*actin:MYL9L-GFP*) embryos at 40 % epiboly (from **Movie S7**). Notice the vegetalward movement of cells and E-YSL (red top arrows) and the retrograde animalward cortical flow from the yolk cell vegetal pole sinking at the E-YSL (red bottom arrows). Scale bar 25 μm. **D**) E-YSL contraction is delayed in Rab5ab YMOs. Between 4.5 to 7.5 hours post fertilization, the E-YSL width is reduced from 80 to 10 μm as an average in controls (blue), but from 80 to 40 μm in Rab5ab YMOs (red). Data collected form surface projections of membrane-GFP (Tg (β-*actin:m-GFP*) *rab5ab* and sibling YMOs (see **Movie S8**). **E)** Compared to control siblings at 70 % epiboly (C YMO - left), leading EVL cells of Rab5ab YMOs flatten and elongate latitudinally (right). Actin was stained with phalloidin-TRITC. Scale bars 25 μm. All confocal images are maximum projections. Quantification of the shape ratio (animal to vegetal vs. latitudinal - Y axis) of leading EVL cells (far right) show that control siblings (red) are extremely elongated on the animal vegetal direction while leading EVL cells in Rab5ab YMOs (blue) flatten at their front and elongate latitudinally. Standard deviations are shown.

The autonomous effects on the yolk cell cortical actomyosin dynamics were accompanied by non-autonomous changes in the shape of EVL cells. EVL cells flattening and elongation (animal to vegetal - AV), which are known outcomes of the accumulation of actin and activated myosin in the E-YSL [26], were prevented in Rab5ab YMOs, with the leading EVL cells elongating in the dorsal to ventral (DV) direction (Fig. 3E and **Movie S8**). We reasoned that these modified shapes respond to alterations in tension anisotropy within the E-YSL, which as a rule increases as epiboly progresses in normal conditions [25].

Last, we found that the stereotyped movements of yolk granules, which are known to passively respond to the cortical stresses created at the E-YSL [25], became disturbed upon Rab5ab depletion. Velocity fields, estimated by Particle Image Velocimetry (PIV) from meridional multiphoton microscopy sections, showed that the yolk granules regular toroidal vortices associated with epiboly [25] were severely affected (see Fig. 4A and **Movie S9**). They became uncoordinated showing a noticeably slower kinetics.

**Figure 4.**
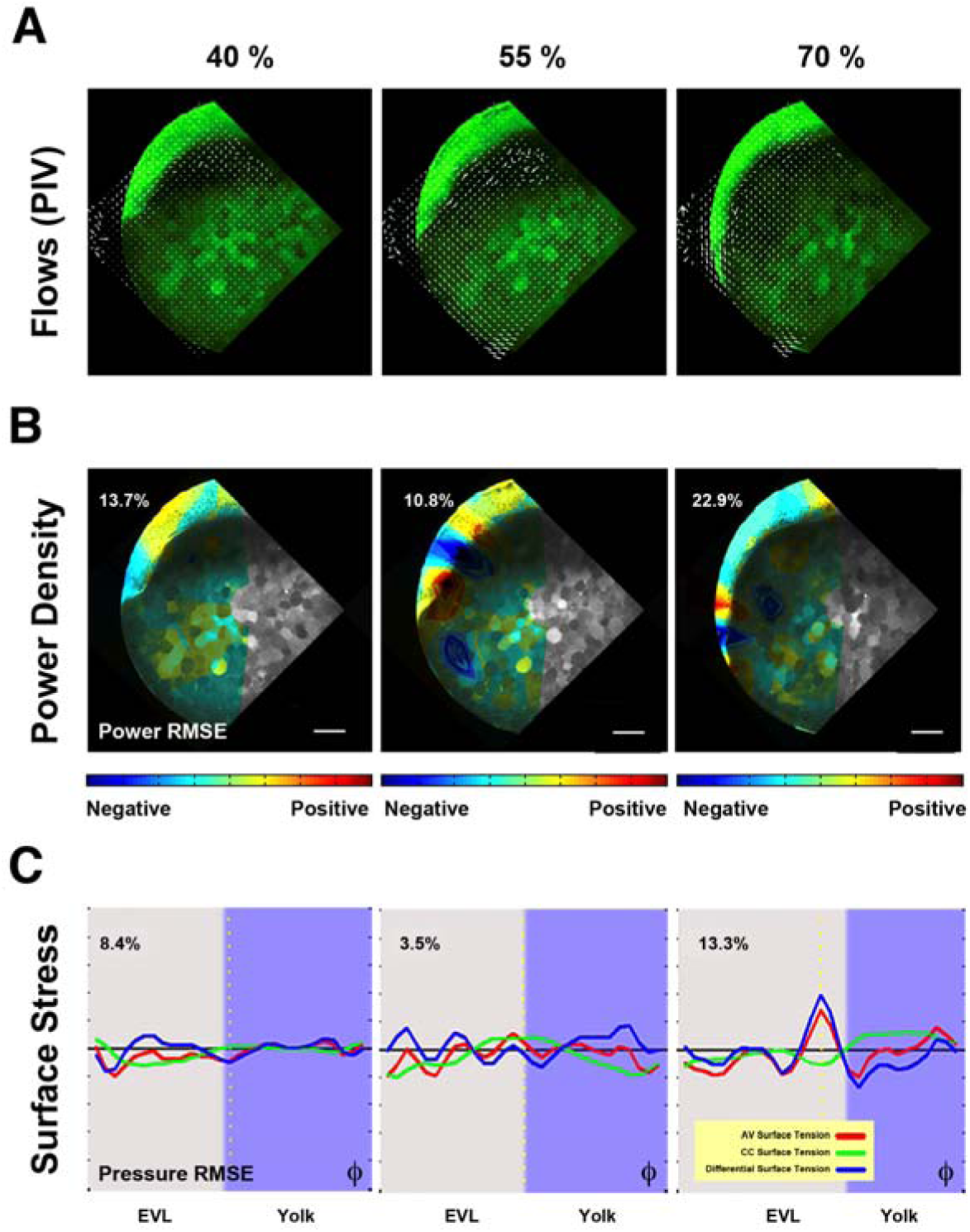
Biomechanics of yolk cell endocytosis impaired embryos. **A**) Yolk granules flows patterns are altered in Rab5ab YMOs. PIV of time-lapse snapshots imaged by two-photon microscopy of a Tg (β-*actin: m-GFP*) Rab5ab YMO (from **Movie S9**; see also **Movie S10**). From epiboly onset, yolk granules flows are uncoordinated in Rab5ab YMOs. The internal toroidal vortices characteristic of epiboly progression [25] do not form or are severely reduced and the laminar flows become asymmetrically distributed. Animal (A) and vegetal (V) poles are indicated. Scale bar 25 μm. **B**) Mechanical power density maps over time obtained by HR analysis in Rab5ab YMOs (from **Movie S11**). The Relative Mean Square Error (RMSE) of the inferred power maps is shown as a percentage expressing the fitting accuracy. Qualitatively, Rab5ab YMOs display no differences with wild type embryos in the spatial and temporal distribution of mechanical work [25] but exhibit an overall decrease of power of about four fold. **C**) Longitudinal along the AV axis (red) and latitudinal or circumferential - CC (green) stresses and their differences (blue) along the embryo cortex in a membrane-GFP transgenic (Tg (β-*actin:m-GFP*)) Rab5ab YMO (from **Movie S12**). Stresses were plotted as a function of the φ angle from animal to vegetal (40 %; 50 %; 65 % epiboly). The equator - dotted yellow line -, yolk cell surface - purple shadow - and the RMSE of the dynamic pressure (fitting accuracy) as a percentage for each time point are displayed. The latitudinal stress does not steep up from animal to vegetal until 60 % epiboly, while the longitudinal stress oscillations are sustained.

In summary, our data indicate that membrane removal at the E-YSL, mainly mediated by Rab5ab, is necessary for the correct structural organization and activity of the EYSL. Endocytic activity in the yolk cell, ahead of the EVL, would influence E-YSL contractility, which in turn would non-autonomously affect both EVL cells elongation and the pattern of yolk granules flows.

### Epiboly mechanics in Rab5ab YMOs

Remarkably, in addition to the structural and functional defects observed upon epiboly arrest as a result of the impairment of yolk cell membrane removal, the overall geometry of Rab5ab YMOs was affected. The final shape of the embryos became rather an ellipsoid than a sphere with an elongated animal to vegetal (AV) axis (see Fig. 2 and **Movie S5**). This elongated shape resulted from both the animalward expansion of the blastoderm after 50 % epiboly and the slight elongation of the yolk cell towards the vegetal pole. The altered geometry of these embryos suggested that their global biomechanics may be compromised and that the topography and dynamics of stresses during epiboly were disrupted in Rab5ab YMOs. To verify this proposal, we determined the spatio-temporal profile of mechanical power and cortical tension of Rab5ab YMOs by Hydrodynamic Regression (HR) [25]. In wild-type embryos throughout epiboly, HR reveals a stereotyped mechanical power density pattern. At the epiboly onset, the active structures mainly map to the blastoderm. Then, once the EVL front crosses the equator, the largest mechanical power density localizes in the active, actomyosin-rich, E-YSL while the adjacent EVL cells opposes deformation. As a consequence, a gradient of tension pointing towards the vegetal pole progressively develops at the yolk cell surface [25].

To define the biomechanical make up of Rab5ab YMOs we employed experimental 2D velocity fields obtained by PIV from time-lapse imaging of meridional sections (see above, Fig. 4A and **Movie S9**). These analyses showed that the yolk granules’ flows were severely impaired (see **Movie S10**). Simulated 3D velocity fields generated from a spherical cortex model were fitted to the experimental velocity fields and dynamic pressure, mechanical power densities and surface tension maps were all inferred by HR [25]. These analyses uncovered an overall four-fold reduction of mechanical energy in Rab5ab YMOs, although the topographical distribution of mechanical power was indistinguishable (Fig. 4B and **Movie S11**) from that of wild-type embryos [25]. Conversely, both longitudinal (AV) and latitudinal (CC) surface stresses and the positive vegetalward gradient of tension in the yolk cell surface of wild-type siblings [25] were significantly decreased (see Fig. 4C and **Movie S12**). These data indicate that Rab5ab-mediated yolk cell endocytosis does not influence where and when mechanical power builds up during epiboly but it is necessary to reach a proper level of cortical tension.

To corroborate the differential tensional topology of the yolk cell cortex inferred by HR in Rab5ab YMOs we employed laser microsurgery [36]. Incisional cuts of the yolk cell cortex result in its immediate recoil with an exponentially decaying speed proportional to its tensional level before ablation [37]. We performed laser cuts at the E-YSL in the AV direction on both Rab5ab YMOs and controls. These analyses confirmed that impeding membrane removal leads to a reduction of surface tension (Fig. 5A). Next, by performing laser cuts parallel to the EVL on the yolk cell cortex at different distances from the margin, we, as inferred by HR, observed that the yolk cell vegetalward gradient of tension characteristic of wild type animals did not develop in Rab5ab YMOs. The averaged recoil velocities for a series of 20 μm laser cuts performed at 65 % epiboly at 20 μm (n = 21) and at 60 μm (n = 9) ahead of the EVL margin showed no statistically significant differences. Conversely, in control wild-type embryos, stress tension (recoil velocity) for equivalent cuts increased by 60 % in the vegetal direction [25].

**Figure 5.**
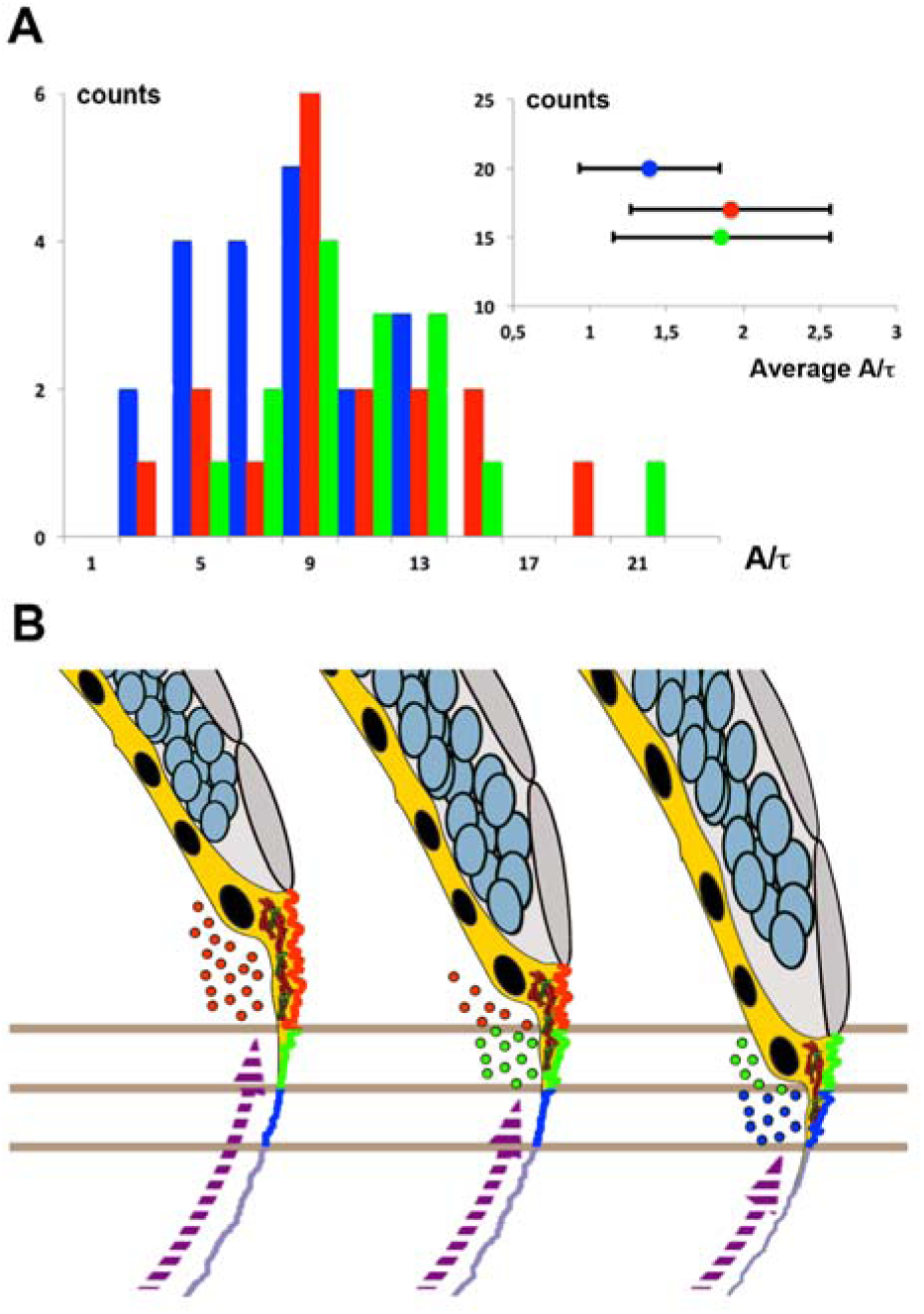
Membrane cortical tension and endocytosis at the E-YSL are necessary for epiboly progression. **A)** The distribution (counts) of instant retraction velocities (A/t) after laser surgery of the actomyosin cortex of Myosin-GFP (Tg (β-*actin:MYL9L-GFP*) Rab5ab YMOs at 55% epiboly (blue) shows a significant reduction (Wilcoxon test p<0.01) versus wild type (red) and control YMOs (green). The instant velocity estimate was extracted from the exponential fit of the distance between fronts (see Supplemental Experimental Procedures). The averaged instant retraction velocity (A/t) for Rab5ab YMOs was 1.39 ± 0.46 μm, while control YMOs reach 1.86 ± 0.71 μm and wild type embryos 1.92 ± 0.65 μm (inset). Counts represent the number of analyzed laser cuts for each condition. **B**) Proposed model of epiboly progression. The contractile E-YSL and the imbalance of stiffness between the EVL and the yolk cell surface account for epiboly progression. Rab5ab-mediated yolk cell localized endocytosis (color coded dots) accounts for the reduction of the yolk cell surface coupled to the progression of EVL (grey) and DCs (blue) towards the vegetal pole. Membrane removal associates to the convolution and contraction of the E-YSL surface and the recruitment of actin and myosin (purple dashed arrow) from vegetally located pools. Three chronological time points are shown. Different sequential zones on the surface of the E-YSL are color-coded. Actin and myosin are diagrammatically illustrated in red and green within the YSL (yellow).

All in all, our data indicate that local membrane removal is essential for strengthening latitudinal (CC) and longitudinal (AV) forces at the E-YSL and for the development of an anisotropic gradient of tension at the cortex. These mechanical elements, jointly, would govern epiboly movements.

## DISCUSSION

The balance between endocytosis and recycling seems to be critical to regulate cell morphology and tissue deformations in multiple morphogenetic processes. Rab5-mediated endocytosis is required downstream of acto-myosin contraction to remove excess membrane in bottle cells in *Xenopus* and to promote their coordinated constriction [6]. Likewise, Rab5 is also required in the amnioserosa during dorsal closure in *Drosophila* for membrane removal as cells delaminate [12]. In zebrafish, different Rab5 isoforms have different roles, participating in Nodal signaling in early embryos [affecting the development of the dorsal organizer (Rab5ab)], or in muscle and brain development (Rab5b and Rab5c respectively) [30]. On the other hand, in fly embryos during cellularization, Rab11, which mediates vesicle recycling, seems to regulate membrane growth and invagination and the elongation of epidermal cells [38].

During epiboly in the zebrafish embryo, the contractile capabilities and gradual change of dimensions of the E-YSL together with the distinct elastic properties of the EVL and the yolk cell surface minimally account for its kinematics and mechanical behavior from 50 % onwards [25]. The E-YSL exerts an isotropic contractile force that generates stress at its adjacent structures, the EVL and the yolk cell cortex, which have different mechanical properties. The EVL is easily deformed by this pulling force and passively expands. On the opposite, the thin yolk cell cortex cannot stretch in response to contraction. In this scenario, we propose that the localized yolk cell membrane removal is essential for the effective movement of the EVL towards the vegetal pole (Fig. 5B). As a result, the non-convoluted yolk cytoplasmic layer would be progressively recruited to the adjacent E-YSL as this is eliminated.

Throughout epiboly, to keep the yolk cell surface balanced, the membrane removal at the front of the advancing EVL must be compensated by membrane recruitment at the I-YSL. Membrane turnover compensation (exocytosis) in the yolk cell remains to be explored, yet, the coupling of exocytosis and endocytosis in EVL cells observed during epiboly [39] provides support for such potential outcome.

Endocytosis of the E-YSL surface was previously suggested to contribute to epiboly [7, 28]. However, knocking-down Dynamin 2 dependent membrane endocytosis in the yolk cell had little effect on epiboly progression [31]. This suggested that the endocytic removal of the yolk cell membrane was dispensable for epiboly. Yet, Dynamin 2 activity only accounted for a fraction of the yolk cell membrane endocytosis and we found that a more efficient inhibition of membrane removal achieved by depleting Rab5ab in the yolk cell led to epiboly arrest. These results point to macropinocytosis and not to clathrin-mediated Dynamin 2-dependent endocytosis as the main mechanism involved in membrane trafficking in the E-YSL. Macropinocytosis is characterized by large non-selective membrane internalization and by the presence of actin cytoskeleton protrusions (ruffles) and has been previously proposed as a plausible for membrane remodeling [2-4]. Indeed, forced macropinocytosis elicited by injection of human Rab5a mRNA in one-cell stage embryos robustly accelerate epiboly progression [40].

A common underlying feature of endocytic membranes, as opposed to other regions of the plasma membrane, is their high curvature [10]. This curvature is somehow linked to the presence of a specific set of regulatory proteins, many of them necessary for curvature generation [41]. Indeed, membrane curvature is influenced by cytoskeleton motor proteins such as myosin [42], which gears pushing (e.g., to get neck membranes closer together) and pulling forces (e.g., to keep vesicle necks under tension) [43]. In this scenario, our data suggest that actomyosin contractility is necessary for the buckling of the E-YSL membrane.

In the absence of Rab5ab in the yolk cell, the overall power and the longitudinal and latitudinal stresses (and the shear stress) at the embryo cortex were severely disturbed. The gradient of tension along the φ axis of the yolk cell surface was extremely weakened (Fig. 4C) and the yolk cell cortical tension suffered a significant reduction (Fig. 5A). Yet, significant actomyosin contractile capability persisted in Rab5ab YMOs and actin and myosin remained localized in the E-YSL cortex. Thus, we propose the existance of a positive regulatory loop in which an initial burst of E-YSL actomyosin contractility is followed by yolk cell membrane endocytosis, which itself potentiates localized cortex constriction. Still, the underlying mechanics of this macropinocytosis-dependent mechanism of actomyosin contraction enhancement in the E-YSL driving epiboly is not clear. Hypotetically, an initial actomyosin contraction at the E-YLS could tense the neighbouring membranes and mechanotransductively activate endocytosis-independent actomyosin contraction leading to its self-induced propagation. In this case, the inhibition of macropinocytosis in Rab5ab YMOs would lead to a higher reservoir of buckled membrane, which would prevent the actomyosin dependent increase of membrane tension and the propagation of such mechanostransductively activation of actomyosin. The general decrease of cortical tension observed in Rab5ab YMOs points into this direction. Mechanotransductive cascades leading to myosin apical stabilization have been indeed described during germ band extension and collective mesoderm constriction in the *Drosophila* embryos [44-46].

Mounting evidence points to a direct relation between membrane reservoir and trafficking pathways with tension in the regulation of cell shape changes and movements in morphogenetic processes [8, 47-53]. During morphogenesis, as tissues change their shapes and sizes, cell membranes dynamically change their area, composition and links to the cortex. As a consequence, membrane tension is subjected to constant modulation [54, 55]. How membrane tension is integrated with the cell’s overall mechanical properties is unknown. In teleosts, pioneering studies in loach uncovered a direct correlation between surface membrane buckles and endocytic-rich domains in early eggs. Further, the experimental decrease of loach eggs surface tension by volume reduction was found to lead to buckling of their membrane [56]. Alongside, in *Fundulus heteroclitus* embryos, mechanical deformations affect epithelial apical membrane turnover [57]. Yet, these early studies failed to provide a comprehensive view of the links between membrane removal, tension and morphogenetic movements.

We propose that, in the early zebrafish embryo, the surface membrane tension constitutes a mechanical buffering system constantly maintained by endocytosis and contractile activity at the E-YSL that regulates epiboly progression. The rates of removal of E-YSL membrane would vary with time and would be proportional to the tension of the yolk cell surface. Endocytosis will lead to membrane tension anisotropies in the yolk cell surface and these will mechanically feedback to regulate membrane dynamics and cortex contractility. This mechanical loop alongside the concerted actions of latitudinal and longitudinal forces at the E-YSL would direct epiboly movements.

It has recently been reported that Rab5 controls a diverse set of collective movements by promoting directional locomotion. In this scenario, multicellular cohorts change their mechanical properties in response to membrane trafficking [40]. Mechanical loops set up by membrane remodeling could constitute a common way to coordinate tissue movements in morphogenetic processes.

## EXPERIMENTAL PROCEDURES

### Zebrafish lines maintenance

AB and TL wild type strains were used throughout this study. Membrane-GFP transgenic (Tg (β-*actin:m-GFP*)) fish [58] were provided by Lilianna Solnica-Krezel and Myosin-GFP transgenics (Tg (β-*actin:MYL9L-GFP*)) (unpublished) by Carl-Philipp Heisenberg. Adult fish were cultured under standard conditions and staged embryos were maintained at 28.5 °C in embryo medium [59].

### mRNA and Morpholino injections

A DNA construct encoding for LifeAct-GFP [60] and cloned in a Zebrafish expression vector was provided by Erez Raz. mRNA was *in vitro* synthesized (μMessage Machine kit, Ambion) and injected into the yolk at one- or 512-cell stages (150 pg). To knockdown Rab5ab, morpholino yolk injections (4 ng and 8 ng) were performed at the 512-1024-cell stage.

### Actin, myosin and nuclear staining

Zebrafish embryos were fixed overnight in 4 % paraformadehyde at 4 °C, washed in 0.3 % Triton in PBS (PBT) and manually dechorionated. They were then washed in PBT, followed by a 2-hour incubation in blocking solution (1 % bovine serum albumin in PBT). Embryos were then incubated either for 1 hour in blocking solution containing 0.2 μg/μl Phalloidin-TRITC (Molecular Probes, Invitrogen) at room temperature or with an antibody against activated myosin (rabbit anti-phospho-myosin light chain 2 Ser19 (Cell Signaling) at 1:100). DAPI was used for nuclear counterstaining. After incubation, embryos were washed 4 times for 15 minutes in PBT. Immunostained embryos were incubated with a secondary fluorescent antibody and washed in PBT. For imaging, embryos were mounted on dishes with 0.5 % low melting agarose (A9045 Sigma) in PBS medium. Images were acquired on a Zeiss LSM700 confocal microscope with 10 X / 0.3 and 63 X / 1.40 oil objectives.

### Live Imaging and Analysis

Whole embryo images were collected from non-dechorionated animals aligned in a 1.2 % agarose mold and covered by E3 medium. Images were acquired (4X magnification) every 5 minutes with an Olympus MVX10 Macroscope.

For confocal and spinning-disc microscopy, embryos were mounted in 0.5 % low melting agarose (A9045 Sigma) in E3 embryo medium.

Sagital sections (350 μm depth from the yolk cell membrane surface) were collected from (Tg (β-*actin: m-GFP*)) embryos using a Leica SP5 two-photon microscope equipped with a mode-locked near-infrared MAITAI Laser (Spectra-Physics) tuned at 900 nm, with non-descanned detectors and with a 25 X / 0.95 water-dipping objective. Images were scanned at 200 Hz and frames were averaged three times. Stacks of 30 μm, 10 μm step-size, were acquired every 2 minutes.

Dextran and lectin internalization were monitored from dechorionated embryos previously incubated in 0.05% 10.000 MW Rhodamine B-Dextran (Life Technologies) for 10 minutes [32] or 100 μg/ml lectin-TRITC (Sigma L1261) for 5 minutes at the sphere stage, both diluted in E3 embryo medium. The Lectin-TRITC used was from *Helix pomatia* [57], which binds *N*-acetyl-D-galactosamine and *N*-acetyl-Dglucosamine residues of glycoproteins and glycolipids on the cell surface. After treatment embryos were rinsed in E3 medium, mounted in 0.5 % low melting agarose and imaged in a Zeiss LSM700 confocal microscope with a 40 X / 1.3 oil immersion objective. A stack of 20 μm, 0.39 μm step size, was acquired every 4 minutes.

To visualize myosin cortical flows, spinning-disc images were captured from (Tg (β-*actin: MYL9L-GFP*)) embryos on either an Olympus X81 inverted microscope (Andor Technologies), using a 40 X / 0.60 Dry objective or a Zeiss Axiovert 200M inverted microscope (PerkinElmer UltraView ERS) using a 40 X / 1.3 oil DIC objective. Stacks of 16 μm, step size 1 μm, were acquired every 45 seconds.

To visualize the surface of the yolk cell, (Tg (β-*actin: m-GFP*)) embryos were imaged in a Zeiss LSM700 confocal microscope with a 63 X / 1.4 oil objective. A stack of 25 μm, step size of 0.2 μm was acquired. We also used embryos injected with *LifeAct-GFP* at the 512-cell stage and collected the images with a Zeiss Axiovert 200M inverted microscope (PerkinElmer UltraView ERS) using a 100 X / 1.4 oil DIC objective. Stacks of 10μm, step size of 0.45 μm, were acquired every 12 seconds.

For photo-bleaching, selected ROIs were created for lectin-TRITC soaked embryos and bleached using 100 % power of a 555 nm laser with 100 iterations in the selected area (in the YSL at 150 μm from the EVL margin) in embryos at 40 % epiboly. A stack of 4 μm, step size 1 μm, was acquired every 30 seconds.

Most image analyses and processing were performed using Fiji (<u>http://pacific.mpicbg.de</u>) and Matlab (Mathworks). To obtain velocity fields we applied the MatPIV software package written by Johan Kristian Sveen for use with Matlab [61]. To measure the width of the wrinkled area, surface projections at different stages were obtained with Matlab and mean width and standard deviations were plotted (Excel, MS Office). To quantify endocytosis, E-YSL dextran-containing vesicles were monitored from maximum projections of Z-stack images.

Laser Surgery Experiments, Retraction Analysis and Hydrodynamic Regression (HR) are described in the Supplemental Experimental Procedures.

## ACKNOWLEDGEMENTS

We thank the Confocal Microscopy Unit from IBMB-PCB, the Advanced Digital Microscopy Core Facility from IRB Barcelona, Xavier Esteban and members of the laboratory for continuous support. We are grateful to Lila Solnica-Krezel and Carolina Minguillón for reading earlier versions of this manuscript. The Consolidated Groups Program of the Generalitat de Catalunya and DGI and Consolider Grants from the Ministry of Economy and Competitivity of Spain to EMB supported this work.

## AUTHOR CONTRIBUTION

MM and AHV performed all biological tests; PAP designed the modeling and the regression analysis and EMB designed the study, analyzed the data and wrote the paper. MM and AHV contributed equally to the study. All authors discussed the results and commented on the manuscript.

## SUPPLEMENTAL EXPERIMENTAL PROCEDURES AND INFORMATION

Supplemental Experimental Procedures

Supplementary Figure Legends

- Figure S1

Supplementary Movies Legends

- Movie S1, Related to Figure 1
- Movie S2, Related to Figure 1
- Movie S3, Related to Figure 2
- Movie S4, Related to Figure 2
- Movie S5, Related to Figure 2
- Movie S6, Related to Figure 3
- Movie S7, Related to Figure 3
- Movie S8, Related to Figure 3
- Movie S9, Related to Figure 4
- Movie S10, Related to Figure 4
- Movie S11, Related to Figure 4
- Movie S12, Related to Figure 4

## SUPPLEMENTAL EXPERIMENTAL PROCEDURES

### Morpholinos

Morpholinos (MOs) were purchased from Gene Tools and designed against selected regions (ATG or UTR) of the *rab5ab* gene (Accession Number ENSDARG00000007257) (Gene Tools): MO1-ATG (5-TCGTTGCTCCACCTCTTCCTGCCAT-3), MO2-UTR (5-GACCCAAAACCCCAATCTCCTGTAC-3), MO3-UTR-ATG (5-ACCTCTTCCTGCCATGACCCAAAAC-3) and a mismatch MO (5-TCcTTcCTCgACCTCTTCgTcCCAT-3) (mispaired nucleotides in lower case). Interference with *rab5c* (ENSDARG00000026712)] was performed with the following MO: 5-CGCTGGTC-CACCTCGCCCCGCCATG-3 provided by C.P. Heisenberg [34]. For all experiments, a group of embryos was injected with a Standard Control MO (5-CCTCTTACCTCAGTTACAATTTATA-3).

### Hydrodynamics Regression (HR)

HR is based in fitting analytically modeled velocity fields to experimental velocity fields in and outside a cortex. Considering that in deforming tissues, stresses at the fluid/cortex boundary are continuous (boundary condition), HR can estimate cortical stresses and retrieve the complete dynamic pressure distribution in the fluid and at the fluid-cortex interface. From these, HR also infers at each time point the cortex shear stress at each point of the surface and the mechanical power density. HR is performed independently at each time point to retrieve the overall spatio-temporal distribution of all these mechanical quantities [24].

In our analyses, experimental 2D velocity fields were estimated by PIV from time-lapse imaging of meridional sections of zebrafish embryos. Second, simulated 3D velocity fields generated from a spherical cortex model (SC) with Stokeslets pairs distributed on a single spherical shell were fitted to the experimental velocity fields. Last, knowing the fluid deformation rates, it is possible to calculate the local values of the cortical surface tension, the cortical mechanical power density and the spatio-temporal evolution of both cortical stresses and mechanical power density maps (analytical codes are available on [24].

### Laser Surgery Experiments and Retraction Analysis

Laser surgery of the actomyosin cortex was performed with a pulsed UV laser (355 nm, 470 ps per pulse) by inducing plasma-mediated ablation as described before [36]. To compare the cortical tension in the longitudinal direction at the E-YSL a 20 μm-laser line containing 50 pulses was scanned 5 times at a frequency of 500 Hz, parallel to the EVL front, centering the cut at a distance of about 20 μm, through a 63 X / 1.2 W objective lens. Fluorescence imaging was performed through a custom spinning Nipkow disc unit equipped with a 488 nm laser line and a Hamamatsu ORCA CCD camera, acquiring at 1.5 frames per second. Transmission and fluorescence imaging was performed by alternated illumination with two out-of-phase mechanical shutters blocking the 488 nm laser and the halogen bright field lamp.

We followed the accepted assumption [36] that the tension present in the actomyosin cortex before the laser cut is proportional to the outward velocity of the immediate recoil. Retraction analysis was performed through a customized kymograph analysis along the retraction axis (perpendicular to the cut), with ImageJ (http://rsbweb.nih.gov/ij/). Kymograph processing included subtraction of the intensity minimum and normalization to the maximum, both measured in the position of the cut, to ensure stable edge detection by intensity threshold across the whole sequence. The front-to-front length, during the retraction phase (until reaching a plateau), was fitted to an exponential function (Igor Pro 6.0, Wavemetrics) to evaluate the slope at the origin.

The function used was:

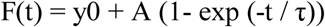

and the slope at the origin was derived from its derivative:

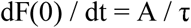

The width of photo bleaching (about 1 μm) introduced by the UV laser was subtracted to the measured length L. This method was applied to the comparative analysis of YMOs conditions.

## SUPPLEMENTAL FIGURE

**Figure S1.**
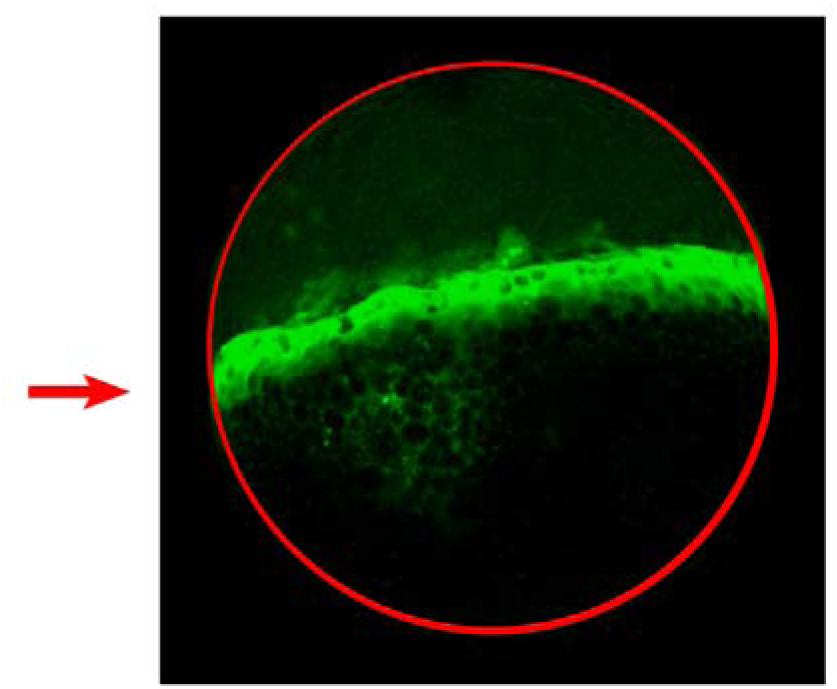
Yolk Cell-Specific Morphants (YMOs) Fluorescent image of an embryo injected at the 512 cells stage with a fluorescently labeled Morpholino (FITC) observed at 40 % Epiboly. The red arrow points to the fluorescence in the yolk cell, accumulating at the E-YSL.

## SUPPLEMENTAL MOVIES LEGENDS

**Movie S1. Endocytosis**

Confocal video time-lapse of a wild type animal soaked in lectin–TRITC for 5 minutes at sphere stage. 20 μm width stacks were captured every 4 minutes. Note the cumulative lectin-TRITC internalization at the E-YSL domain ahead of the advancing EVL.

**Movie S2. Endocytosis at the E-YSL**

HiLo LUT code (Fiji) processed confocal video time-lapse of a lectin-TRITC soaked embryo showing a circular photo bleached ROI in the yolk cell 60 μm away from the EVL leading edge. 4 μm width stacks were captured every 30 seconds. Maximum projection is shown. Scale bar 25 μm. The photo-bleached membrane (blue circle) is not removed or endocytosed and remains unchanged up to its enclosure within the advancing E-YSL.

**Movie S3. Endocytosis impairment in Rab5ab YMOs**

Confocal video time-lapse of control and Rab5ab YMOs (medium dose) soaked in Rhodamine B-Dextran for 5 minutes at sphere stage. 20 μm width stacks were captured every 4 minutes. Note the reduced dextran internalization ahead of the advancing EVL.

**Movie S4. Specific arrest of epiboly movements in Rab5ab YMOs**

Confocal video time-lapse of a membrane-GFP (Tg (β-*actin:m-GFP*) Rab5ab YMOs (medium dose) that slowly proceeds throughout epiboly arresting at 70 % presenting an open back phenotype but succeeding in other gastrulation movements leading to somite formation.

**Movie S5. Medium dose Rab5ab YMO macroscopic phenotype**

Macroscope video time-lapse of control and Rab5ab YMO sibling embryos (at a middle dose) are shown. Bright field images were captured every 5 minutes from sphere stage to 16 hours post fertilization. Note that the Rab5ab YMO at a middle dose is at 60 % epiboly at the time when the control embryo closes (yellow arrow). The Rab5ab YMO continues its elongation in the animal-vegetal direction but remains open.

**Movie S6. High dose Rab5ab YMO macroscopic phenotype**

Macroscope video time-lapse of control and Rab5ab YMO sibling embryos (at a high dose) are shown. Bright field images were captured every 5 minutes from sphere stage to 16 hours post fertilization. Note that Rab5ab YMO at a high dose halt before reaching the equator and burst.

**Movie S7. Cortical Myosin flows on the E-YSL in control versus Rab5ab YMOs**

Spinning-disc video time-lapses of zebrafish transgenic (Tg (β-*actin:MYL9L-GFP*)) embryos from 50% epiboly onwards (control sibling (top) and a Rab5ab YMO (bottom). The movements of cortical actin were analyzed by Particle Image Velocimetry (PIV). 9 μm width stacks were captured every 12 seconds. Maximum projections are shown. Scale bar 25 μm. The retrograde animalward cortical myosin flows from the vegetal pole of the yolk cell towards the E-YSL are impaired in Rab5ab YMOs.

**Movie S8. EVL marginal cells elongate in the CC direction in Rab5ab YMOs**

Confocal video time-lapse of control and *rab5ab* yolk cell membrane labeled [Tg (*β-actin:m-GFP*)] zebrafish transgenic YMOs (medium dose). While control embryos marginal EVL cells elongate in the animal/vegetal direction (left), Rab5ab YMOs marginal cells flatten and elongate in the circumferential direction. The speed of progression of the EVL is reduced and this eventually stalls.

**Movie S9. Yolk granules flows in Rab5ab YMOs**

Flow trajectories from PIV measurements of a two-photon video time-lapse of a membrane-GFP (Tg (β-*actin:m-GFP*)) zebrafish transgenic Rab5ab YMO at the middle plane, 350 μm deep inside, throughout epiboly. Scale bar 100 μm. Flows in the Rab5ab YMO are partly arrested.

**Movie S10. Yolk granules kinematics in Rab5ab YMOs**

Flow lines from PIV measurements of two-photon excitation stitched video time-lapses of membrane-GFP (Tg (β-*actin:m-GFP*)) zebrafish transgenic control (left) and *rab5ab* (right) YMO siblings at the middle plane, 350 μm deep inside, throughout epiboly. Scale bar 100 μm. The Rab5ab YMO shows epiboly delay and disrupted yolk flows.

**Movie S11. Power Density Maps in Rab5ab YMOs**

Two-photon excitation video time-lapse of a membrane-GFP (Tg (β-*actin: m-GFP*)) Rab5ab YMO at the middle plane, 350 μm deep inside, throughout epiboly (left). Mechanical power density maps were calculated by HR from the velocity fields, color-coded at an equal scale for each time point (right) and overlaid on the fluorescence images (negative - blue; 0 - green; positive - red) (middle). The relative mean square error for the Power is displayed as a percentage for each time point. Scale bar 100 μm. As in the wild type [24], during doming, most of the energy supply is generated by the blastoderm. After 50 % epiboly, the E-YSL constriction becomes the main motor of epiboly, while the EVL cells resist their stretching. Thus, Rab5ab YMOs display no qualitative differences with wild type embryos in their mechanical activity distribution but, importantly, they exhibit an overall decrease of power of about four fold.

**Movie S12. Longitudinal and latitudinal stresses in Rab5ab YMOs**

Main surface stresses were calculated by HR of velocity fields obtained from stitched two-photon excitation video time-lapse of a membrane-GFP (Tg (β-*actin:m-GFP*)) Rab5ab YMO embryo at the middle plane, 350 μm deep inside, throughout epiboly. Longitudinal (red) and latitudinal (green) stresses and their differences (blue) were plotted as a function of the φ angle from animal to vegetal. Time points were every 40 minutes. The equator - dotted yellow line -, yolk cell surface - purple shadow - and the relative mean square error of the pressure as a percentage for each time point are displayed. In these YMOs, the longitudinal and latitudinal stresses are equal at the poles. Contrary to wild type embryos, the latitudinal stress does not steep up from animal to vegetal. Their difference (longitudinal minus latitudinal) shows a defined profile at the EVL margin after 50 % epiboly, becoming positive in the EVL and negative at the E-YSL.

## REFERENCES

1. McMahon, H.T., and Boucrot, E. (2011). Molecular mechanism and physiological functions of clathrin-mediated endocytosis. Nat Rev Mol Cell Biol 12, 517–533.

2. Lim, J.P., and Gleeson, P.A. (2011). Macropinocytosis: an endocytic pathway for internalising large gulps. Immunol Cell Biol 89, 836–843.

3. Cao, H., Chen, J., Awoniyi, M., Henley, J.R., and McNiven, M.A. (2007). Dynamin 2 mediates fluid-phase micropinocytosis in epithelial cells. J Cell Sci 120, 4167–4177.

4. Swanson, J.A., and Watts, C. (1995). Macropinocytosis. Trends in cell biology 5, 424–428.

5. Lecuit, T., and Pilot, F. (2003). Developmental control of cell morphogenesis: a focus on membrane growth. Nat Cell Biol 5, 103–108.

6. Lee, J.Y., and Harland, R.M. (2010). Endocytosis is required for efficient apical constriction during Xenopus gastrulation. Current biology: CB 20, 253–258.

7. Betchaku, T., and Trinkaus, J.P. (1986). Programmed endocytosis during epiboly of Fundulus heteroclitus. American zoologist 26, 7.

8. Gauthier, N.C., Masters, T.A., and Sheetz, M.P. (2012). Mechanical feedback between membrane tension and dynamics. Trends in cell biology 22, 527–535.

9. Satoh, D., Sato, D., Tsuyama, T., Saito, M., Ohkura, H., Rolls, M.M., Ishikawa, F., and Uemura, T. (2008). Spatial control of branching within dendritic arbors by dynein-dependent transport of Rab5-endosomes. Nat Cell Biol 10, 1164–1171.

10. Doherty, G.J., and McMahon, H.T. (2009). Mechanisms of endocytosis. Annu Rev Biochem 78, 857–902.

11. Fabrowski, P., Necakov, A.S., Mumbauer, S., Loeser, E., Reversi, A., Streichan, S., Briggs, J.A., and De Renzis, S. (2013). Tubular endocytosis drives remodelling of the apical surface during epithelial morphogenesis in Drosophila. Nature communications 4, 2244.

12. Mateus, A.M., Gorfinkiel, N., Schamberg, S., and Martinez Arias, A. (2011). Endocytic and recycling endosomes modulate cell shape changes and tissue behaviour during morphogenesis in Drosophila. PloS one 6, e18729.

13. Zeigerer, A., Gilleron, J., Bogorad, R.L., Marsico, G., Nonaka, H., Seifert, S., Epstein-Barash, H., Kuchimanchi, S., Peng, C.G., Ruda, V.M., et al. (2012). Rab5 is necessary for the biogenesis of the endolysosomal system in vivo. Nature 485, 465–470.

14. Zerial, M., and McBride, H. (2001). Rab proteins as membrane organizers. Nat Rev Mol Cell Biol 2, 107–117.

15. Barbieri, M.A., Fernandez-Pol, S., Hunker, C., Horazdovsky, B.H., and Stahl, P.D. (2004). Role of rab5 in EGF receptor-mediated signal transduction. Eur J Cell Biol 83, 305–314.

16. Lanzetti, L., Palamidessi, A., Areces, L., Scita, G., and Di Fiore, P.P. (2004). Rab5 is a signalling GTPase involved in actin remodelling by receptor tyrosine kinases. Nature 429, 309–314.

17. Tall, G.G., Barbieri, M.A., Stahl, P.D., and Horazdovsky, B.F. (2001). Ras-activated endocytosis is mediated by the Rab5 guanine nucleotide exchange activity of RIN1. Dev Cell 1, 73–82.

18. Song, S., Eckerle, S., Onichtchouk, D., Marrs, J.A., Nitschke, R., and Driever, W. (2013). Pou5f1-dependent EGF expression controls E-cadherin endocytosis, cell adhesion, and zebrafish epiboly movements. Dev Cell 24, 486–501.

19. Ulrich, F., Krieg, M., Schotz, E.M., Link, V., Castanon, I., Schnabel, V., Taubenberger, A., Mueller, D., Puech, P.H., and Heisenberg, C.P. (2005). Wnt1 1 functions in gastrulation by controlling cell cohesion through Rab5c and E-cadherin. Dev Cell 9, 555–564.

20. Solnica-Krezel, L. (2006). Gastrulation in zebrafish – all just about adhesion? Curr Opin Genet Dev 16, 433–441.

21. Kimmel, C.B., Ballard, W.W., Kimmel, S.R., Ullmann, B., and Schilling, T.F. (1995). Stages of embryonic development of the zebrafish. Developmental dynamics: an official publication of the American Association of Anatomists 203, 253–310.

22. Rohde, L.A., and Heisenberg, C.P. (2007). Zebrafish gastrulation: cell movements, signals, and mechanisms. Int Rev Cytol 261, 159–192.

23. Cheng, J.C., Miller, A.L., and Webb, S.E. (2004). Organization and function of microfilaments during late epiboly in zebrafish embryos. Developmental dynamics: an official publication of the American Association of Anatomists 231, 313–323.

24. Betchaku, T., and Trinkaus, J.P. (1978). Contact relations, surface activity, and cortical microfilaments of marginal cells of the enveloping layer and of the yolk syncytial and yolk cytoplasmic layers of fundulus before and during epiboly. The Journal of experimental zoology 206, 381–426.

25. Hernandez-Vega, A., Marsal, M., Pouille, P.A., Tosi, S., Colombelli, J., Luque, T., Navajas, D., Pagonabarraga, I., and Martin-Blanco, E. (2016). Polarized cortical tension drives zebrafish epiboly movements. EMBO J.

26. Koppen, M., Fernandez, B.G., Carvalho, L., Jacinto, A., and Heisenberg, C.P. (2006). Coordinated cell-shape changes control epithelial movement in zebrafish and Drosophila. Development 133, 2671–2681.

27. Behrndt, M., Salbreux, G., Campinho, P., Hauschild, R., Oswald, F., Roensch, J., Grill, S.W., and Heisenberg, C.P. (2012). Forces driving epithelial spreading in zebrafish gastrulation. Science 338, 257–260.

28. Solnica-Krezel, L., and Driever, W. (1994). Microtubule arrays of the zebrafish yolk cell: organization and function during epiboly. Development *120*, 2443- 2455.

29. Schepis, A., Sepich, D., and Nelson, W.J. (2012). alphaE-catenin regulates cell-cell adhesion and membrane blebbing during zebrafish epiboly. Development 139, 537–546.

30. Kenyon, E.J., Campos, I., Bull, J.C., Williams, P.H., Stemple, D.L., and Clark, M.D. (2015). Zebrafish Rab5 proteins and a role for Rab5ab in nodal signalling. Dev Biol 397, 212–224.

31. Lepage, S.E., Tada, M., and Bruce, A.E. (2014). Zebrafish Dynamin is required for maintenance of enveloping layer integrity and the progression of epiboly. Developmental biology 385, 52–66.

32. Feng, B., Schwarz, H., and Jesuthasan, S. (2002). Furrow-specific endocytosis during cytokinesis of zebrafish blastomeres. Exp Cell Res 279, 14–20.

33. Kimmel, C.B., and Law, R.D. (1985). Cell lineage of zebrafish blastomeres. II. Formation of the yolk syncytial layer. Dev Biol 108, 86–93.

34. Thisse, B., Heyer, V., Lux, A., Alunni, V., Degrave, A., Seiliez, I., Kirchner, J., Parkhill, J.P., and Thisse, C. (2004). Spatial and temporal expression of the zebrafish genome by large-scale in situ hybridization screening. Methods Cell Biol 77, 505–519.

35. Ulrich, F., and Heisenberg, C.P. (2008). Probing E-cadherin endocytosis by morpholino-mediated Rab5 knockdown in zebrafish. Methods Mol Biol 440, 371–387.

36. Colombelli, J., Besser, A., Kress, H., Reynaud, E.G., Girard, P., Caussinus, E., Haselmann, U., Small, J.V., Schwarz, U.S., and Stelzer, E.H. (2009). Mechanosensing in actin stress fibers revealed by a close correlation between force and protein localization. J Cell Sci 122, 1665–1679.

37. Grill, S.W. (2011). Growing up is stressful: biophysical laws of morphogenesis. Curr Opin Genet Dev 21, 647–652.

38. Pelissier, A., Chauvin, J.P., and Lecuit, T. (2003). Trafficking through Rab 11 endosomes is required for cellularization during Drosophila embryogenesis. Current biology: CB 13, 1848–1857.

39. Ahn, H.J., Park, Y., Kim, S., Park, H.C., Seo, S.K., Yeo, S.Y., and Geum, D. (2010). The expression profile and function of Satb2 in zebrafish embryonic development. Molecules and cells 30, 377–382.

40. Malinverno, C., Corallino, S., Giavazzi, F., Bergert, M., Li, Q., Leoni, M., Disanza, A., Frittoli, E., Oldani, A., Martini, E., et al. (2017). Endocytic reawakening of motility in jammed epithelia. Nat Mater.

41. Kozlov, M.M., Campelo, F., Liska, N., Chernomordik, L.V., Marrink, S.J., and McMahon, H.T. (2014). Mechanisms shaping cell membranes. Curr Opin Cell Biol 29, 53–60.

42. Spudich, G., Chibalina, M.V., Au, J.S., Arden, S.D., Buss, F., and Kendrick-Jones, J. (2007). Myosin VI targeting to clathrin-coated structures and dimerization is mediated by binding to Disabled-2 and PtdIns(4,5)P2. Nat Cell Biol 9, 176–183.

43. Roux, A., Uyhazi, K., Frost, A., and De Camilli, P. (2006). GTP-dependent twisting of dynamin implicates constriction and tension in membrane fission. Nature 441, 528–531.

44. Pouille, P.A., Ahmadi, P., Brunet, A.C., and Farge, E. (2009). Mechanical signals trigger Myosin II redistribution and mesoderm invagination in Drosophila embryos. Sci Signal 2, ra16.

45. Fernandez-Gonzalez, R., Simoes Sde, M., Roper, J.C., Eaton, S., and Zallen, J.A. (2009). Myosin II dynamics are regulated by tension in intercalating cells. Dev Cell 17, 736–743.

46. Mitrossilis, D., Roper, J.C., Le Roy, D., Driquez, B., Michel, A., Menager, C., Shaw, G., Le Denmat, S., Ranno, L., Dumas-Bouchiat, F., et al. (2017). Mechanotransductive cascade of Myo-II-dependent mesoderm and endoderm invaginations in embryo gastrulation. Nature communications 8, 13883.

47. Kremnyov, S.V., Troshina, T.G., and Beloussov, L.V. (2012). Active reinforcement of externally imposed folding in amphibians embryonic tissues. Mech Dev 129, 51–60.

48. Diz-Munoz, A., Fletcher, D.A., and Weiner, O.D. (2013). Use the force: membrane tension as an organizer of cell shape and motility. Trends in cell biology 23, 47–53.

49. Apodaca, G. (2002). Modulation of membrane traffic by mechanical stimuli. American journal of physiology. Renal physiology 282, F179–190.

50. Gauthier, N.C., Fardin, M.A., Roca-Cusachs, P., and Sheetz, M.P. (2011). Temporary increase in plasma membrane tension coordinates the activation of exocytosis and contraction during cell spreading. Proceedings of the National Academy of Sciences of the United States of America 108, 14467–14472.

51. Dai, J., and Sheetz, M.P. (1995). Regulation of endocytosis, exocytosis, and shape by membrane tension. Cold Spring Harbor symposia on quantitative biology 60, 567–571.

52. Dai, J., Sheetz, M.P., Wan, X., and Morris, C.E. (1998). Membrane tension in swelling and shrinking molluscan neurons. The Journal of neuroscience: the official journal of the Society for Neuroscience 18, 6681–6692.

53. Sheetz, M.P., and Dai, J. (1996). Modulation of membrane dynamics and cell motility by membrane tension. Trends in cell biology 6, 85–89.

54. Figard, L., and Sokac, A.M. (2014). A membrane reservoir at the cell surface: unfolding the plasma membrane to fuel cell shape change. Bioarchitecture 4, 39–46.

55. Clark, A.G., Wartlick, O., Salbreux, G., and Paluch, E.K. (2014). Stresses at the cell surface during animal cell morphogenesis. Current biology: CB 24, R484–494.

56. Ivanenkov, V.V., Meshcheryakov, V.N., and Martynova, L.E. (1990). Surface polarization in loach eggs and two-cell embryos: correlations between surface relief, endocytosis and cortex contractility. The International journal of developmental biology 34, 337–349.

57. Fink, R.D., and Cooper, M.S. (1996). Apical membrane turnover is accelerated near cell-cell contacts in an embryonic epithelium. Dev Biol 174, 180–189.

58. Cooper, M.S., Szeto, D.P., Sommers-Herivel, G., Topczewski, J., Solnica-Krezel, L., Kang, H.C., Johnson, I., and Kimelman, D. (2005). Visualizing morphogenesis in transgenic zebrafish embryos using BODIPY TR methyl ester dye as a vital counterstain for GFP. Dev Dyn 232, 359–368.

59. Westerfield, M. (2000). The Zebrafish Book: A Guide for the Laboratory Use of Zebrafish (Danio rerio), (Eugene, OR: University of Oregon Press).

60. Riedl, J., Crevenna, A.H., Kessenbrock, K., Yu, J.H., Neukirchen, D., Bista, M., Bradke, F., Jenne, D., Holak, T.A., Werb, Z., et al. (2008). Lifeact: a versatile marker to visualize F-actin. Nat Methods 5, 605–607.

61. Supatto, W., Debarre, D., Moulia, B., Brouzes, E., Martin, J.L., Farge, E., and Beaurepaire, E. (2005). In vivo modulation of morphogenetic movements in Drosophila embryos with femtosecond laser pulses. Proceedings of the National Academy of Sciences of the United States of America 102, 1047–1052.

